# Discovering Novel Cell Types across Heterogeneous Single-cell Experiments

**DOI:** 10.1101/2020.02.25.960302

**Authors:** Maria Brbić, Marinka Zitnik, Sheng Wang, Angela O. Pisco, Russ B. Altman, Spyros Darmanis, Jure Leskovec

## Abstract

Although tremendous effort has been put into cell type annotation and classification, identification of previously uncharacterized cell types in heterogeneous single-cell RNA-seq data remains a challenge. Here we present MARS, a meta-learning approach for identifying and annotating known as well as novel cell types. MARS overcomes the heterogeneity of cell types by transferring latent cell representations across multiple datasets. MARS uses deep learning to learn a cell embedding function as well as a set of landmarks in the cell embedding space. The method annotates cells by probabilistically defining a cell type based on nearest landmarks in the embedding space. MARS has a unique ability to discover cell types that have never been seen before and annotate experiments that are yet unannotated. We apply MARS to a large aging cell atlas of 23 tissues covering the life span of a mouse. MARS accurately identifies cell types, even when it has never seen them before. Further, the method automatically generates interpretable names for novel cell types. Remarkably, MARS estimates meaningful cell-type-specific signatures of aging and visualizes them as trajectories reflecting temporal relationships of cells in a tissue.

## Introduction

High-throughput single-cell transcriptional profiling has enabled remarkable progress in our understanding of cellular mechanisms of the disease and development [1–4]. Cell atlas datasets, including Mouse Cell Atlas [5,6] and Human Cell Atlas [7], systematically measure the transcriptome of individual cells in multiple sites in the organism and at several time points during growth and development. These datasets have contributed to the discovery of novel cell types and cell transcriptional states [8–11]. However, to assist with the identification of new cell types, there is currently a big gap as it requires techniques that (1) harmonize heterogeneous and time-varying datasets, (2) learn dataset-invariant cell representations, and (3) use the learned representations in order to decide whether groups of measured cells represent previously uncharacterized cell types and cell states. Such techniques would have the power to reveal novel cell types, enable investigation of biology that underlies those cell types and their cellular activity, and would thus form a crucial tool in an expanding single-cell computational toolbox.

Existing single-cell tools train deep neural network models to learn how to embed cells into a vector space. Importantly, the structure of the space is optimized during model training to reflect geometry of the training dataset [12–17]. After the method learns cell embeddings, it clusters them to find groups of cells with similar gene expression programs. Finally, the method then annotates/assigns each group to a cell type for which enough annotated cells already exist in the training dataset [18, 19]. However, present methods are unable to annotate cells that are not characterized in existing datasets or have not been measured before. Present methods cannot classify cells into new cell types that do not exist in the training data. While recent semi-supervised and supervised methods [20–23] have made initial steps towards empowering single-cell analyses by reusing previously annotated datasets, these methods require that all cell types have many annotated examples in the training data. As a result, current methods are unable to identify novel/unseen cell types.

Here we introduce MARS, an approach for annotating known/seen as well as novel/unseen cell types in heterogeneous and time-varying single-cell datasets. MARS uses meta-learning, a paradigm in machine learning that focuses on efficient use of limited annotations [24–27]. In particular, MARS first constructs a meta-dataset by integrating (i) any number of single-cell experiments in which cells are annotated (*i.e.*, labeled) by a cell type, and (ii) an unannotated experiment, which does not necessarily share any cell types with the labeled data. Using the meta-dataset, MARS jointly learns a set of cell-type landmarks and an embedding function that projects cells into a shared embedding space, such that cells are close to their cell-type landmarks. The embedding space, learned by a deep neural network, identifies gene expression programs and leverages commonalities between experiments in the meta-dataset. This gives MARS a unique ability to generalize to unannotated experiments and identify cell types that were never seen during training. We apply MARS to Tabula Muris [6] and Tabula Muris Senis [28] cell atlases. We find that MARS successfully transfers knowledge between diverse tissues and aligns the same cell types, even when they originate from different tissues. Further, we find that MARS learns meaningful cell-type-specific signatures of aging in a mouse. Our results show that MARS considerably outperforms current techniques for cell type classification. Remarkably, MARS is able to accurately identify cell types it has never seen during training and can probabilistically recommend interpretable names for them.

## Results

### Meta-learning in MARS

MARS takes as input single-cell gene expression profiles from heterogeneous or time-varying experiments, such as different tissues or stages of development. MARS creates a meta-dataset that consists of (i) experiments in which cells are annotated according to their cell types, and (ii) a completely unannotated experiment in which cell types are unknown. The unannotated experiment can originate from different source and does not need to share any cell types with the annotated experiments. The goal then is to annotate cells in the unannotated experiment, such as never-before-seen tissue or stage of development. This is a novel setup not considered by previous single-cell methods.

### Overview of MARS

Given a meta-dataset as input, MARS learns a set of cell-type landmarks and a non-linear embedding function. The embedding function projects a high-dimensional expression profile of each cell to a low-dimensional vector (*i.e.*, cell embedding), that directly captures the cell-type identity (Fig. 1a). Cell-type landmarks are defined as cell type representatives and are learned for both annotated and unannotated experiments. The embedding function is a deep neural network that maps cells to the embedding space. The embedding space is defined such that cells embed close to their cell-type landmarks. The embedding function is shared between all experiments in the meta-dataset, which gives MARS the ability to generalize to an unannotated experiment and to capture the similarity of cell types across annotated and unannotated experiments.

**Figure 1:**
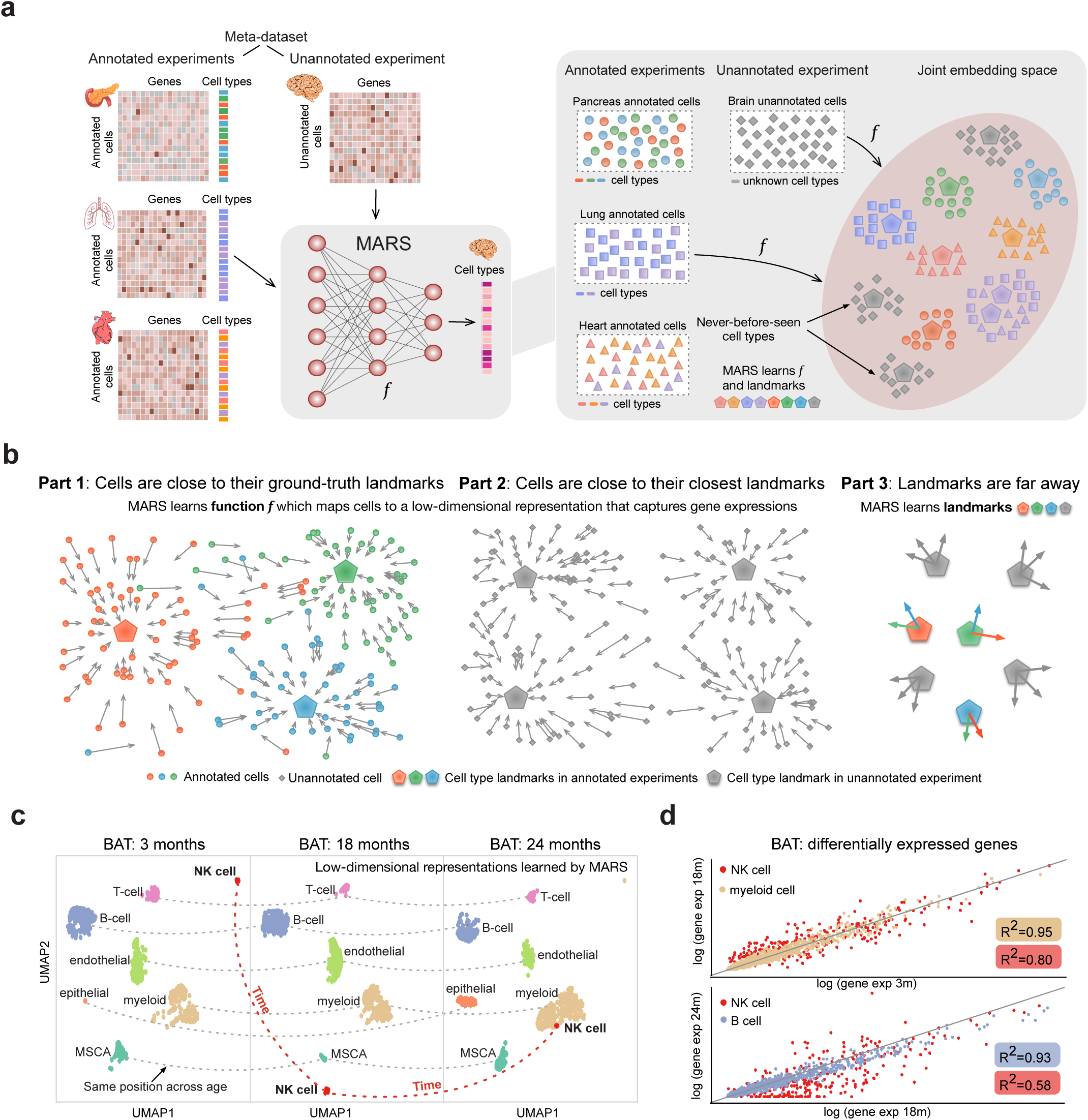
MARS is a meta-learning approach for discovery of novel cell types across heterogeneous single-cell experiments. **(a)** Illustration of the MARS method. Given a set of heterogeneous annotated experiments (e.g. pancreas, lung, heart tissues), MARS aims to annotate a new, completely unannotated experiment (e.g. brain tissue), even if it does not have any cell type in common with annotated experiments. Using deep neural networks, MARS projects all cells in the meta-dataset (annotated and unannotated) to the shared embedding space and learns nonlinear embedding function *f* such that cells from the same cell types are embedded close to each other, while cells from different cell types are embedded far away. **(b)** MARS relies on the notion of a cell type landmarks. Objective function of MARS simultaneously optimizes three parts: (i) within annotated experiment, distance to the ground-truth landmark is minimized; (ii) within unannotated experiment, distance to the closest landmark is minimized; and (iii) within each experiment, distance between landmarks is maximized. Cell type landmarks and experiment-invariant cell representations are learned jointly and in an end-to-end fashion. **(c)** MARS reconstructs a trajectory of brown adipose tissue (BAT) cell types during the life span of a mouse. All BAT cell types except natural killer (NK) cells retain the same position across three different time points, while the effect of aging is reflected on NK cells, implying that their gene expression profiles change over time. **(d)** Comparison of gene expression of differentially expressed genes in BAT across different time points (Benjamini-Hochberg FDR adjusted p-value < 0.1; t-test). Top plot shows average gene expression of differentially expressed genes of 3 months and 18 months old mouse for NK cells and myeloid cells. Bottom plot shows average gene expression of 18 months and 24 months old mouse for NK cells and B-cells. Variability in gene expressions of NK cells is higher than in other cell types, indicating that MARS detects biologically meaningful aging patterns.

Mathematically, MARS uses regularization in the form of pretraining the neural network with a deep autoencoder that minimizes a data reconstruction error (Methods). The pretraining step serves as a prior towards configuration of the parameter space useful for the generalization to a novel unannotated dataset. Using the pretrained network as initialization, MARS then learns mapping of all cells to the shared embedding space such that similar cells are close to each other, while dissimilar cells are far way. Equipped with the concept of cell-type landmarks, we design an objective function that aims to learn a representation in which cells group close to their corresponding landmarks (Methods). The objective function consists of three parts (Fig. 1b): (i) in the annotated experiments, the distance between cell embeddings and ground truth cell-type landmark is minimized; (ii) in the unannotated experiment, the distance between cell embeddings and the nearest cell-type landmark is minimized; and (iii) distance between cell-type landmarks within each experiment is maximized. The rationale is to encourage cells from the same cell type to have similar representations, while representations of cells from different cell types are far apart. MARS does not impose any constraint on the radius of a discovered cell type so cell types can form clusters that reflect their transcriptional similarity to other cell types.

### MARS identifies cell-type-specific signatures of aging

We assess MARS’s ability to infer cell-type trajectories on the Tabula Muris Senis dataset [28], covering the life span of a mouse. In particular, we analyze whether the same cell types from different time points are embedded close together (*i.e.*, aligned) in the embedding space. We use the brain adipose tissue (BAT) data from 3 months, 18 months, and 24 months old mice as the annotated experiments. We find that MARS aligns all cell types except the set of natural killer (NK) cells. NK cells change their position at every time point (Fig. 1c), indicating the existence of transcriptional changes. To confirm that the motion of NK cells as detected by MARS is meaningful, we further analyze the variability in gene expression of differentially expressed genes across three time points. Populations of NK cells indeed show higher variability than other cell types with a coefficient of determination (*R*^2^) of 0.79 between 3 months and 18 months old mice, and 0.58 between 18 months and 24 months old mice (Fig. 1d). In contrast, the median of *R*^2^ of other cell types is 0.93 (Q1-Q3: 0.89 − 0.95) and 0.89 (Q1-Q3: 0.84 − 0.89), respectively. Furthermore, populations of NK cells share 6% of differentially expressed genes across three time points compared to the average of 26.8% shared genes on other cell types in brown adipose tissue, confirming that the representation learned by MARS captures transcriptional changes in aging NK cells. Moreover, this finding has been well-characterized experimentally [29–31], suggesting that cellular functions of NK cells are impaired in aging mice and can lower the resistance to cancer and pathogenic microorganisms.

### MARS outperforms other methods by a large margin

To demonstrate the performance of MARS on a cell type annotation task, we use the manually curated Tabula Muris dataset [6]. We consider each tissue as a separate experiment (Methods and Supplementary Note 1). We leave one tissue out as unannotated and use all others as annotated experiments. We then test the performance on the unannotated held-out tissue experiment. Note that most often the unannotated held-out tissue shares no cell-types with the annotated tissues, which means that MARS has to be able to identify completely new cell types it has never seen during training.

We compare MARS to four methods that can also apply to this task: deep generative model ScVi [12], kernel-learning approach SIMLR [32], and two community detection approaches Leiden [33] and Louvain [34], which are employed in two widely used single-cell analysis tools, including Scanpy [35] and Seurat [36] (Supplementary Note 2). Remarkably, MARS achieves 31% better adjusted Rand index score than the second-best performing SIMLR (Fig. 2a). When measuring performance using various metrics, including accuracy, adjusted mutual information score, and F1 score, MARS retains significantly better performance than all other methods (Supplementary Fig. 1). Of note, MARS uses the same set of parameters across all tissues and shows high robustness to the choice of the neural network architecture. In particular, MARS’s performance is not affected even when the embedding dimension changes (Supplementary Fig. 2).

**Figure 2:**
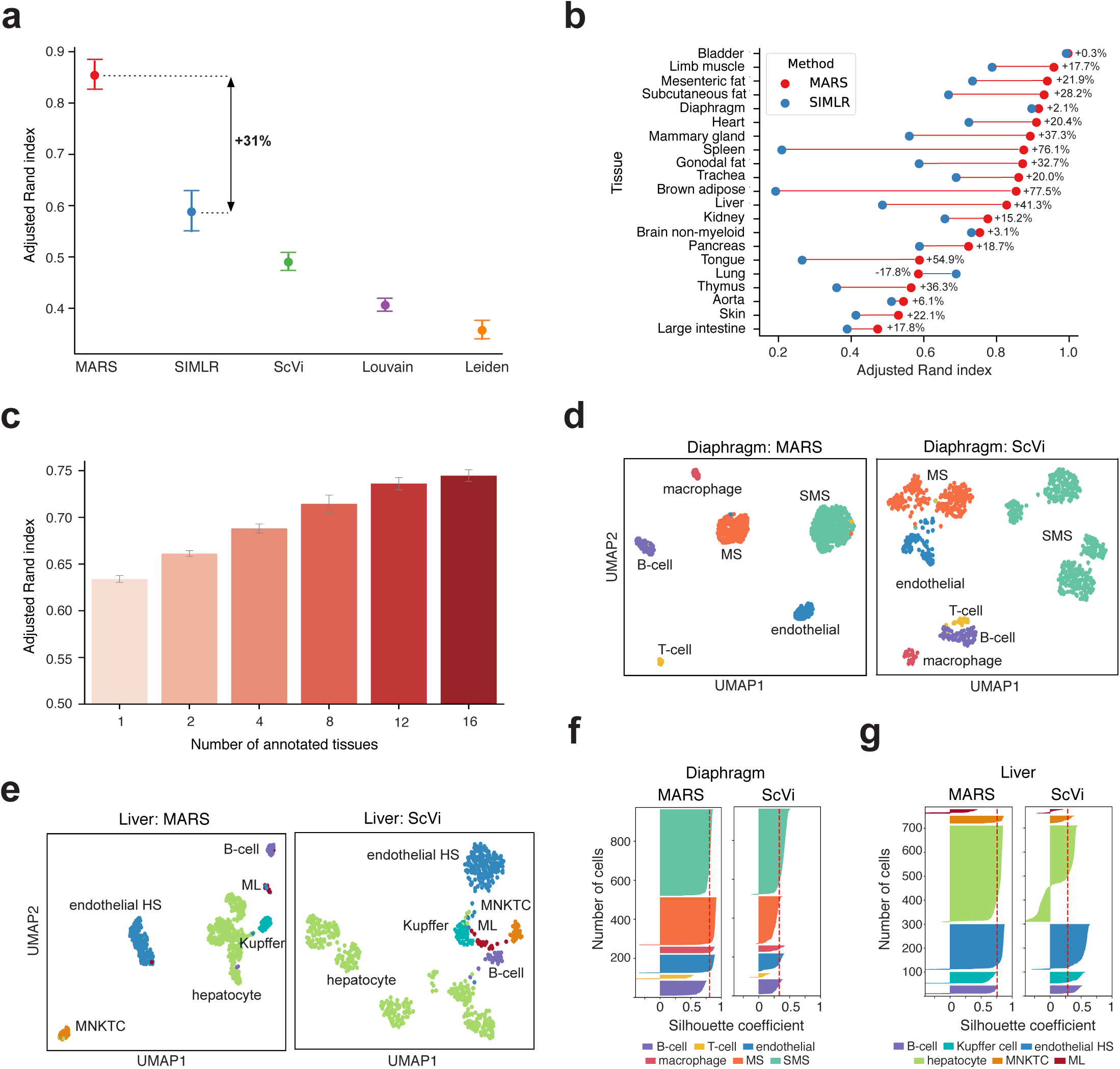
MARS achieves positive learning transfer and accurately annotates cells. Performance is evaluated on the Tabula Muris dataset [6] (*n* = 44,516 annotated cells). **(a)** Median performance of MARS and four baseline methods evaluated using adjusted Rand index (ARI) score across 21 different tissues (Methods). Higher value indicates better performance, where 1.0 is perfect performance and 0.0 indicates random clustering. Error bars are standard errors estimated by bootstrapping cells within tissue with n=20 iterations. MARS is trained in leave-one-tissue-out manner, and the held out tissue was completely unannotated (see Methods). MARS improves the score of the second best performing SIMLR [32] by 31.1%. **(b)** Comparison of the MARS’s performance on individual tissues with the SIMLR. Tissues are ranked according to the MARS’s ARI score. MARS significantly outperforms SIMLR (*p* = 1*e*^*−*4^; Wilcoxon signed-rank test), and achieves better performance on all except one tissue. **(c)** Effect of the number of annotated tissues in the meta-dataset on MARS’s performance. Bars show average adjusted Rand index and standard deviation across 20 runs of the method. Annotated tissues are selected based on their similarity to an unannotated tissue, where similarity is computed as the euclidean distance of the mean gene expressions profiles between tissues. MARS improves performance when more tissues are included in the meta-dataset, implying that cross-tissue positive knowledge transfer is achieved even when tissues do not have similar gene expressions. **(d, e)** UMAP visualizations of deep variational autoencoder ScVi’s and MARS’s embeddings for **(d)** diaphragm tissue, and **(e)** liver tissue. SMS stands for skeletal muscle cell, MS for mesenchymal stem, HS for hepatic sinusoid, and MNKTC for mature NK T-cell. Color indicates Tabula Muris cell type annotation. Only cell types with more than five annotated cells are shown. In the MARS’s embedding space, cell types naturally form clusters that correspond to cell types, agreeing well with Tabula Muris annotations. In contrast, the distinction between clusters is not clear in the ScVi’s embedding space and different cell types are often mixed. **(f, g)** Quality of the neural embeddings of MARS and ScVi measured as silhouette coefficient on **(f)** diaphragm tissue, and **(g)** liver tissue. The silhouette score measures mean intra-cluster distance and the mean inter-cluster distance for each data point, indicating how well is a data point matched to its own cluster. The measure ranges between −1 and 1, where 1 corresponds to perfect score. On both tissues, the silhouette coefficient score of MARS is 0.8, whereas ScVi achieves 0.3 score.

We also evaluate performance of MARS on individual tissues, each consisting of many cell types. Compared to the second best performing method SIMLR, MARS performs better on 20 out of 21 tissues (Fig. 2b), and achieves 26.1% higher area under the curve than SIMLR, and 30.1% compared to ScVi (Supplementary Fig. 3). In particular, for heart tissue which contains 7 out of 11 never-before-seen cell types, MARS improves SIMLR’s ARI score by 20.5%. In addition to the tissue-level performance, we also evaluate cell type-level performance between MARS and SIMLR, where we observe that MARS performs especially well on small cell-types with very few cells, cell types with very few differentially expressed genes, and cell types it has never seen during training (Supplementary Fig. 4, 5).

### MARS achieves positive knowledge transfer across tissues

We also show that MARS achieves better performance as the number of the annotated experiments increases. Specifically, we start with the meta-dataset consisting of only one annotated experiment, and then gradually add more annotated experiments in the meta-dataset (Methods). We find that MARS performs considerably better on large meta-datasets (Fig. 2c). In particular, when using heart and mesenteric fat as the unannotated experiments, MARS improves by 35.3% and 30.4%, respectively (Supplementary Fig. 6). Although subcutaneous fat, mesenteric fat, heart and brown adipose tissues do not share any cell types in common with large intestine tissue, including them into meta-dataset when predicting cell types of large intestine improves performance by 10.1%. This analysis demonstrates that MARS effectively reuses annotated experiments, even when they differ in their gene expression profiles from the unannotated experiment. Our results suggest that more annotated experiments yield higher-quality cell embeddings.

### MARS discovers novel cell types and subtypes

We visualize representations of cells learned by MARS in the 2-dimensional UMAP [37] space. MARS learns to embed similar cells close to each other, while dissimilar cells are embedded far, agreeing well with the Tabula Muris annotations. In contrast, in the embedding space learned by ScVi different cell types are often mixed together without a clear decision boundary between cell types (Fig. 2d, e). To quantitatively evaluate the quality of the neural embeddings, we use silhouette coefficient which compares inter- and intracluster distance of data points with −1 as the lowest and 1 as the highest score. On both tissue, MARS achieves silhouette coefficient score of 0.8, whereas ScVi achieves score of 0.3 (Fig. 2f, g).

We also demonstrate that MARS discovers novel cell subtypes. In particular, we analyze mammary gland tissue for which the cell types discovered by MARS differ from the Tabula Muris annotations. MARS separates cells annotated as luminal epithelial cells by Tabula Muris into two different clusters (Fig. 3a). To check whether luminal epithelial cells in two clusters detected by MARS are indeed different, we run a permutation test, comparing Jaccard similarity of Gene Ontology [38] enriched terms of differentially expressed genes in the sampling distribution to Jaccard similarity of clusters detected by MARS (Methods). Results confirm that the gene expression of luminal epithelial cells in clusters detected by MARS differs significantly (Fig. 3b), indicating that MARS discovers subtypes of luminal epithelial cells.

**Figure 3:**
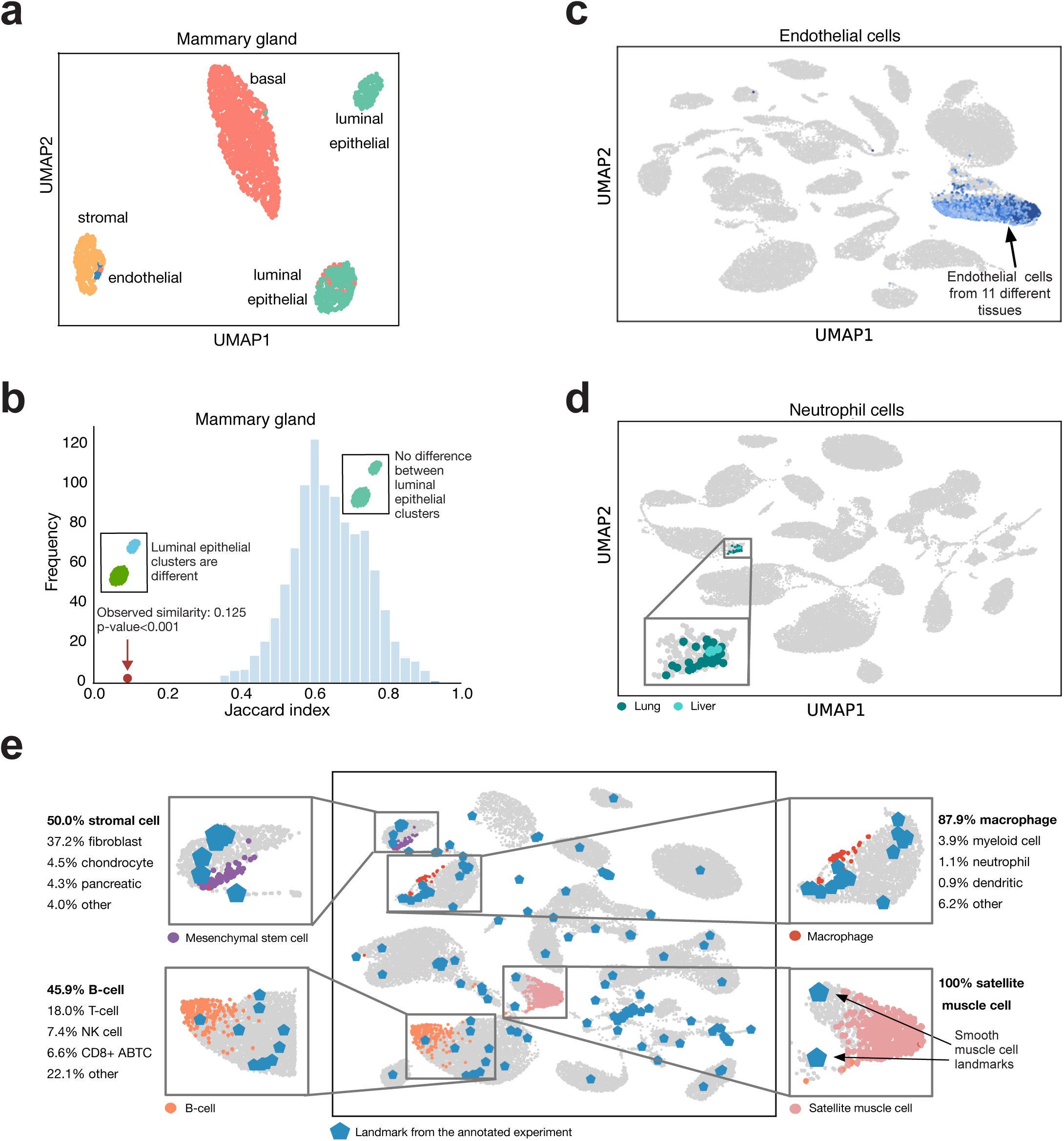
MARS accurately identifies cell types, even when tissues have no cell types in common, and automatically generates interpretable names for novel cell types. **(a)** UMAP visualization of MARS’s embedding of mammary gland tissue cells. MARS indicates that cells annotated as luminal epithelial cells by Tabula Muris annotations belong to two separate clusters. **(b)** Results of permutation test under the null hypothesis that there is no difference between luminal epithelial cells. We define test statistic to be Jaccard similarity of enriched gene ontology terms of differentially expressed genes (Benjamini-Hochberg FDR adjusted p-value < 0.1; t-test) between two groups (see Methods). Observed value is the similarity between two clusters of luminal epithelial cells found by MARS, while distribution is obtained by randomly permuting luminal epithelial cells into two groups with *n* = 1000 iterations. Observed difference between two clusters found by MARS is significant with *p* < 10^*−*3^, implying that MARS recognizes cell subtypes of luminal epithelial cells. **(c, d)** UMAP visualization of MARS joint embedding space of all tissues. Same cell types across different tissues are extremely well aligned, even cell type consisting of only few cells. **(c)** Endothelial cells are aligned across 11 different tissues (brain non-myeloid, diaphragm, brown adipose tissue, subcutaneous fat, mesenteric fat, gonodal fat, limb muscle, mammary gland, pancreas, trachea and thymus), where thymus is used as unannotated tissue. **(d)** Small clusters of neutrophil cells from lung and liver are aligned, where lung is used as unannotated tissue. **(e)** Overview of the MARS cell type naming approach. For unannotated cell type which we want to name, MARS determines distances to all landmarks from the annotated experiments and for each of them outputs probability that discovered cell type should receive the same name (see Methods). In the example, limb muscle is used as unannotated tissue. MARS accurately assigns names to stromal cells, B-cells, macrophages and satellite muscle cells. PDC stands for plasmacytoid dendritic cell and CD8+ ABTC for CD8-positive alpha-beta T-cell.

### MARS correctly aligns and annotates cell types across tissues

MARS utilizes meta-dataset to learn the embedding, which effectively generalizes to never-before-seen experiments. We next examine whether the same cell types across tissues in the annotated and unannotated experiments are embedded close to each other. Out of 105 different cell types in the Tabula Muris dataset, only 20 cell types are present in two or more tissues. We first investigate the endothelial cells, which appear in 11 tissues, and use thymus tissue as an unannotated experiment. We select thymus because it is the most challenging. We find that endothelial cells are exceptionally well aligned across diverse tissues, even in the challenging thymus tissue (Fig. 3c). We observe near-perfect alignment for other cell types that appear across many tissues, such as B-cells (Supplementary Fig. 7). We further evaluate small neutrophil cell type that appears in only lung and liver tissues by using lung as an unannotated experiment. Remarkably, neutrophils from unannotated lung tissue align well to only four liver neutrophil cells (Fig. 3d). Finally, we note that MARS is complementary to integrative approaches for batch-correction, including [17, 22, 23, 39], and can be applied to batch-corrected datasets returned by those approaches.

### MARS can name new cell types

Last, we demonstrate MARS’s ability to assign interpretable names to discovered groups of cells. MARS relies on the cell-type landmarks in the annotated experiments to probabilistically define cell type based on its region in the low-dimensional embedding space. Probabilities are assigned to landmarks in proportion to their probability density under Gaussian centered at a target unannotated cell type (Methods). To demonstrate our approach, we analyze whether cell types with more than 10 cells from the limb muscle tissue are correctly assigned. Indeed, MARS accurately identifies satellite muscle cells and endothelial cells with 100% probability, macrophages with over 87% probability, and B-cells with more than 45% probability (Fig. 3e). At first glance, it may look like MARS misclassifies mesenchymal stem cells (MSC) by assigning them to stromal cells with a high confidence; however, MSC are adherent stromal cells [40]. Furthermore, with 37.2% of probability MSC are assigned to the fibroblast cell type, that is indistinguishable from MSC using morphology and cell-surface markers [40, 41]. Hence, distances in MARS’s embedding space can also be used to infer similarity between cell types.

## Discussion

MARS has a unique ability to transfer knowledge of cell embeddings across heterogeneous experiments that possibly do not have any cell types in common. In doing so, MARS introduces a practical setting for the analysis of single-cell data, in which the experiment of interest can be completely new and unannotated, thereby requiring generalization to never-before-seen cell types.

MARS addresses this challenge by learning cell-type-specific landmarks and a nonlinear embedding function that maps all cells in a joint low-dimensional embedding space shared by annotated and unannotated experiments. Using the learned landmarks of cell types to identify new cell types, MARS provides a framework for annotation of discovered cell types by probabilistically assigning cell types in the neighborhood of the annotated landmarks. As a result, MARS can considerably alleviate the post-hoc manual analyses of cell types.

MARS allows for knowledge transfer across tissues and time-varying experiments. Our approach has important implications for other types of knowledge transfer, including the transfer of cell type annotations across species and different omics measurements, and transfer of cell states across related diseases.

Finally, MARS is complementary to tools for correcting batch effects and data integrative studies, including Scanorama [39], Harmony [17], and Seurat V3 [22]. Results returned by these tools can be directly used as input to MARS. As new comprehensive atlas datasets are generated in line with Human Cell Atlas [7] efforts, we envision that MARS will become an essential tool to help in unraveling unknown cellular diversity of healthy and diseased tissues.

## Methods

### Dataset processing

We downloaded raw read Tabula Muris [6] and Tabula Muris Senis [28] datasets with cell type annotations (see “Data availability”). We filtered low-quality cells with fewer than 5,000 reads and 500 genes, as well as genes expressed in less than 5 cells. We used Scanpy [35] to normalize each cell to 10,000 read counts, and then log transformed the data. Finally, we scaled the dataset to unit variance and zero mean, and we truncated values with maximum value set to 10. The normalization and scaling steps remove any experiment-specific differences and enable alignment based on relative gene expression values. After preprocessing, the number of retained genes was 22,903. The number of annotated cells was 105,960 in Tabula Muris Senis and 44, 516 in Tabula Muris. The number of cells per dataset ranged from 906 to 13,417 cells in Tabula Muris Senis, and 366 to 5,067 cells in Tabula Muris. To demonstrate the ability of MARS to detect aging signatures, we used Tabula Muris Senis dataset. For all other analyses, we used Tabula Muris dataset with re-annotations from Tabula Muris Senis. Additional details are provided in Supplementary Note 1.

### Overview of MARS

The key idea in the MARS model is that representation that encourages clustering of cells in one experiment, also helps in learning to separate cells in a distinct experiment. We aim to accomplish the goal of learning experiment-invariant representation by transferring knowledge of the right distance metric from previously annotated experiments to a new, completely unannotated experiment. We refer to the set of all experiments (annotated and unannotated) over which MARS learns as a meta-dataset, *i.e.*, dataset for learning to learn representation that can easily adapt to new tasks. To achieve transferable features, MARS learns shared representation across all experiments in the meta-dataset. Specifically, given gene expression profiles and cell type assignments in the annotated experiments, and gene expression profiles of an unannotated target experiment, MARS learns nonlinear mapping function *f*_*θ*_ that maps cells from all experiments into a joint embedding space such that cells are grouped according to their cell types. Function *f* is parameterized by learnable feature mapping parameters *θ* of a deep neural network. MARS consists of two stages: (i) pretraining on unannotated target experiment with deep autoencoder, and (ii) learning cell type landmarks and nonlinear cell embedding with deep neural network. MARS optimizes cell type landmarks and parameters *θ* of the experiment-invariant nonlinear mapping function in an end-to-end manner.

#### 1) Pretraining

We first pretrain MARS with an autoencoder. An autoencoder network takes as input normalized gene expression profiles of unannotated experiment **X**^*u*^ ∈ ℝ^*N* ×*G*^, where *N* denotes number of cells and *G* denotes number of genes. Input is mapped to a lower dimensional dense representation vector (encoding). The decoder part maps encoding vector to the reconstruction of the input 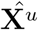. Autoencoder is trained to minimize reconstruction loss 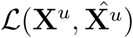, given as the mean squared error between **X**^*u*^ and 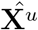. After pretraining, we remove the decoder part and use learned weights to initialize deep network.

#### 2) Initialization of cell-type landmarks

To initialize cell type landmarks, we first map all cells into a lower-dimensional representation vector learned by autoencoder. Then, for each experiment in the meta-dataset we separately run K-means clustering in the embedding space. We use ten random initializations and take the best one in terms of the sum of squared distances of cells to their closest cluster landmark.

#### 3) Loss function

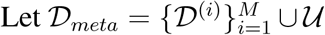 be a set of (*M* + 1) distinct experiments to which we refer to as a meta-dataset. We assume that each experiment 𝒟^(*i*)^ consists of a matrix of normalized gene expression profiles 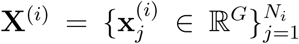, and a vector of cell type annotations 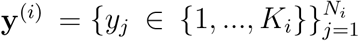, where *G* denotes number of genes, *N*_*i*_ number of cells and *K*_*i*_ number of cell types in a experiment 𝒟^(*i*)^. Furthermore, let 𝒰 consists of a matrix of gene expression profiles 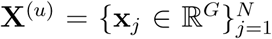 with unknown cell annotations, where *N* denotes number of cells in 𝒰. Given meta-dataset 𝒟_*meta*_, MARS learns cell type landmarks in the annotated experiments 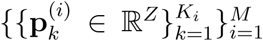, cell type landmarks in the unannotated experiment 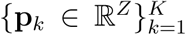, and a nonlinear mapping function *f*_*θ*_ : ℝ^*G*^ → ℝ^*Z*^, where *K* denotes number of cell types in the unannotated experiment, *Z* is dimension of the embedding space, and *θ* are learnable parameters. In MARS, we seek to find joint embedding space such that within each experiment cells group around a single cell type landmark and landmarks are far away. Therefore, the mapping function *f*_*θ*_ is shared between all experiments in the meta-dataset and maps all cells into the joint embedding space.

In the annotated meta-dataset, cell type annotations are known and MARS encourages cells to be close to their ground-truth cell type landmarks. For each annotated experiment 𝒟^(*i*)^ ∈ 𝒟_*meta*_, MARS incorporates the following part in the objective function:

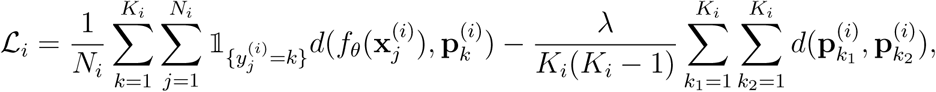

where *λ* is a regularization constant, 𝟙 denotes indicator function, and *d* is distance function. We use squared Euclidean distance as a distance function, but others can be easily incorporated. Of note, all distances are calculated in the low-dimensional embedding space. The first part measures intra-cluster distance between cells and ground-truth landmarks, whereas the second part measures inter-cluster distance between all pairs of landmarks. Intra-cluster distance is minimized to achieve compact representations within a cluster, whereas inter-cluster distance is maximized to push representations of distinct landmarks far away from each other.

Next, we include in the objective function term that encourages clustering structure of the unannotated experiment 𝒰. With the same intuition as above, we again measure intra- and intercluster distance. However, in this case cell type assignments are unknown, so MARS minimizes the distance to the closest cell type landmark in the unannotated experiment. Formally, for 𝒰 ∈ 𝒟_*meta*_ MARS extends the objective function with the following term:

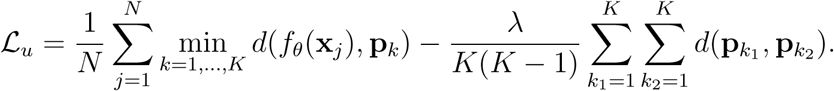

The final objective function optimizes for the annotated and unannotated experiments jointly:

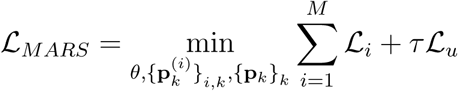

The objective function balances between intra-cluster minimization and inter-cluster maximization. Importantly, both parts are optimized within each experiment, allowing clusters across experiments to align with each other. Cluster landmarks and representation parameters *θ* learned by deep neural network are optimized simultaneously. In each iteration, we first optimize for landmarks while fixing the parameters *θ*. Then, we optimize for *θ* while fixing the landmarks. In the annotated experiments, landmarks are obtained in the closed-form solution. In the unannotated experiment, we update landmarks with the Adam optimizer.

#### 4) Inference

Embeddings of cells in the meta-dataset are obtained by the representation learned in the last layer of the neural network. At the inference time, we annotate cells from the unannotated experiment. In particular, MARS embeds cells from the unannotated experiment into the learned shared embedding space and assigns them to the cluster of the closest cell type landmark.

#### 5) Cell type naming

MARS probabilistically assigns interpretable names to a discovered cluster by relying on the annotated cell type landmarks in the meta-dataset. Probabilities are estimated for every cell type seen in the annotated experiments in proportion to their probability density under Gaussian distribution centered at the mean of a discovered cluster. Then, annotations are assigned to the discovered cluster based on the annotations of the most similar annotated landmarks. Formally, given cell type landmarks 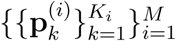 in the annotated experiments, conditional probability that *j*th cluster in the unannotated experiment adds *k*th landmark from the annotated experiments in the set of the most similar landmarks is calculated as follows:

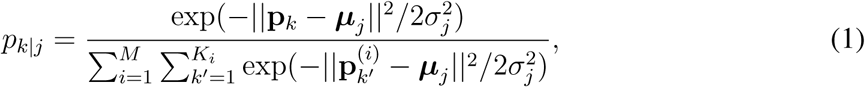

where ***µ***_*j*_ is the mean of cell embedding vectors assigned to target cluster *j*, and *σ*_*j*_ is estimated based on the standard deviation of pairwise Euclidean distances of cells assigned to cluster *j*. Empirically, we observe that embedding data points beforehand in the low-dimensional space with UMAP improves the results. We used 10 UMAP components.

### Implementation and hyper-parameters

In MARS, neural network consists of two encoding and two decoding layers. We used 1000 neurons in the first layer, and 100 neurons in the second layer of the neural network, with a symmetric decoder. We then fine-tuned the parameters with the meta-learning loss introduced in MARS. Best parameters were optimized in the small grid search according to the best mean performance across all tissues. We used Adam optimizer with learning rate 0.001. Activities of the neurons were normalized using layer normalization which estimates the normalization statistics over all hidden units in the same layer. ELU function, defined as *ELU* (*x*) = *max*(0, *x*) + *min*(0, *α*(*exp*(*x*) − 1)), was used as a nonliner activation with *α* set to 1. We pretrained the network for 25 epochs, and fine-tuned for 30 epochs. Regularizer *λ* and *τ* in MARS’s objective function was set to 0.2 and 1, respectively. We assesed robustness of MARS to the selection of architecture by varying embedding dimension across a range of possible values, while keeping all other parameters fixed (Supplementary Fig. 2).

### Performance evaluation

We evaluated MARS performance in leave-one-tissue-out manner. We used all except one tissue as the set of annotated experiments, and held out tissue as an unannotated experiment. We evaluated performance by comparing cell type assignments of the unannotated experiment to the ground-truth clusters. To evaluate how the number of annotated experiments in the meta-dataset affects performance, we used as annotated experiments *n* most similar tissues to unannotated tissue, while varying *n* from 1 to 16. Similarity between tissues was computed as the Euclidean distance of their mean gene expression profiles.

### Visualization

We visualized cell embeddings using UMAP [37]. Cell neighborhood graph was calculated with number of neighbors set to 30. For alignment and annotation visualization, we calculated neighborhood graph and performed UMAP on MARS’s cell embeddings across all tissues.

### Differential gene expression

We performed differential gene expression analysis using Scanpy package. We used t-test as statistical test, and Benjamini-Hochberg method for the adjustment of p-values. Maximum number of genes was set to 1,000.

### Permutation test and functional enrichment analyses

To check whether two clusters of luminal epithelial cells in Fig. 3a are significantly different, we performed permutation test. We chose Jaccard similarity of enriched Gene Ontology (GO) [38] terms between differentially expressed genes of two samples as the test statistic. To calculate differential gene expression, the reference set of cells consisted of all cells that are not annotated as luminal epithelial cells (stromal, basal, and endothelial cells). The observed value of the test statistic was Jaccard similarity of enriched GO terms between differentially expressed genes of two clusters of luminal epithelial cells detected by MARS. Sampling distribution of the test statistic was estimated by randomly permuting luminal epithelial into two groups and calculating Jaccard similarity between the groups. GO [42] enriched terms were calculated using GOATOOLS package [43]. GO terms were propagated to parent terms before functional enrichment tests were calculated. P-values were adjusted using the Benjamini-Hochberg method with FDR< 0.1.

## Data availability

Tabula Muris Senis dataset used in the paper is publicly available from the website at https://figshare.com/projects/Tabula_Muris_Senis/64982. Raw data from the Tabula Muris dataset is publicly available at https://doi.org/10.6084/m9.figshare.5829687.v8. We retrieved data from the website on November 2nd, 2019.

## Code availability

MARS is written in Python using the PyTorch library. The source code is available on Github (https://github.com/snap-stanford/mars).

## Author contribution

M.B., M.Z., and J.L. conceived the study, designed and performed research, contributed new analytic tools, analyzed data, and wrote the manuscript. M.B. also implemented the model, performed experiments, and developed metrics. S.W. discussed the results and contributed to the writing. A.O.P. helped procure and interpret the datasets. S.D. and R.B.A. supervised research and contributed to the writing. J.L. supervised research and the entire project.

## Author information

The authors declare no conflict of interest.

