## Supplementary Information for "Discovering Novel Cell Types across Heterogeneous Single-cell Experiments"

This PDF file includes:

Supplementary Notes 1 and 2

Supplementary Figures 1 to 8

Supplementary References

### Supplementary Note 1 Datasets and preprocessing

We downloaded Tabula Muris Senis [1] datasets with annotations from https://figshare.com/projects/Tabula_Muris_Senis/64982. Raw data for the Tabula Muris dataset is obtained from https://doi.org/10.6084/m9.figshare.5829687.v8. Due to the updated Tabula Muris annotations on Tabula Muris Senis dataset, we used annotations for both datasets from Tabula Muris Senis. In particular, we used field *cell_ontology_class_reannotated* as cell type labels. For cross-age transfer analysis, we used Tabula Muris Senis dataset for 3 months, 18 months and 24 months old mouse. For cross-tissue transfer analysis, we used Tabula Muris dataset. We observed that all methods are incapable to distinguish cell types in brain myeloid tissue that consists of microglial and macrophage cell types. The difficulty is biologically explainable by many shared molecular markers between these cell types, among which TMEM119 is the only known stable marker highly expressed by microglial cells but not expressed by macrophages [2]. Furthermore, microglial cells cover 99% of cells in brain myeloid of Tabula Muris dataset [3], making it hard to detect small macrophage cluster. For that reason, we did not include brain myeloid tissue in the analysis. At the time of writing the paper, marrow tissue annotations were not validated by expert so we did not perform any experiments on the marrow tissue. After filtering brain myeloid and marrow, 21 tissues remained in the dataset. Pretraining of the MARS was performed on the unannotated tissue of the Tabula Muris Senis dataset.

### Supplementary Note 2 Baseline methods

We compared MARS to four unsupervised methods used for clustering single-cell data: Louvain [4], Leiden [5], SIMLR [6] and ScVi [7].

For Louvain and Leiden, we first performed PCA and retained 43 principal components[3]. We computed neighborhood graph with number of neighbors set to 30. We used Scanpy's implementations of Louvain and Leiden with the resolution parameter set to 1.0.

For SIMLR and ScVi, we used implementations provided by the authors. For SIMLR we first performed PCA and retained 500 principal components. Number of neighbors for constructing cell-cell similarity graph was set to 30. For ScVi we first pretrained network with variational autoencoder for 150 epochs. We tried two pretraining strategies: (i) pretraining on the same data as MARS (only unannotated tissues from Tabula Muris Senis), and (ii) pretraining on the Tabula Muris data from all tissues. The latter achieved better performance, so we used that setting in all our analysis and comparison with the ScVi method. After pretraining, we used all parameters recommended by the authors. Specifically, we trained network for 200 epochs with learning rate 0.001 and Adam optimizer. Neural network consisted of two layers with widths 128 and 32. To obtain clustering assignments, ScVi relies on K-means clustering of learned cell embeddings. Since K-means depends on initialization, we reported mean score across 20 runs.

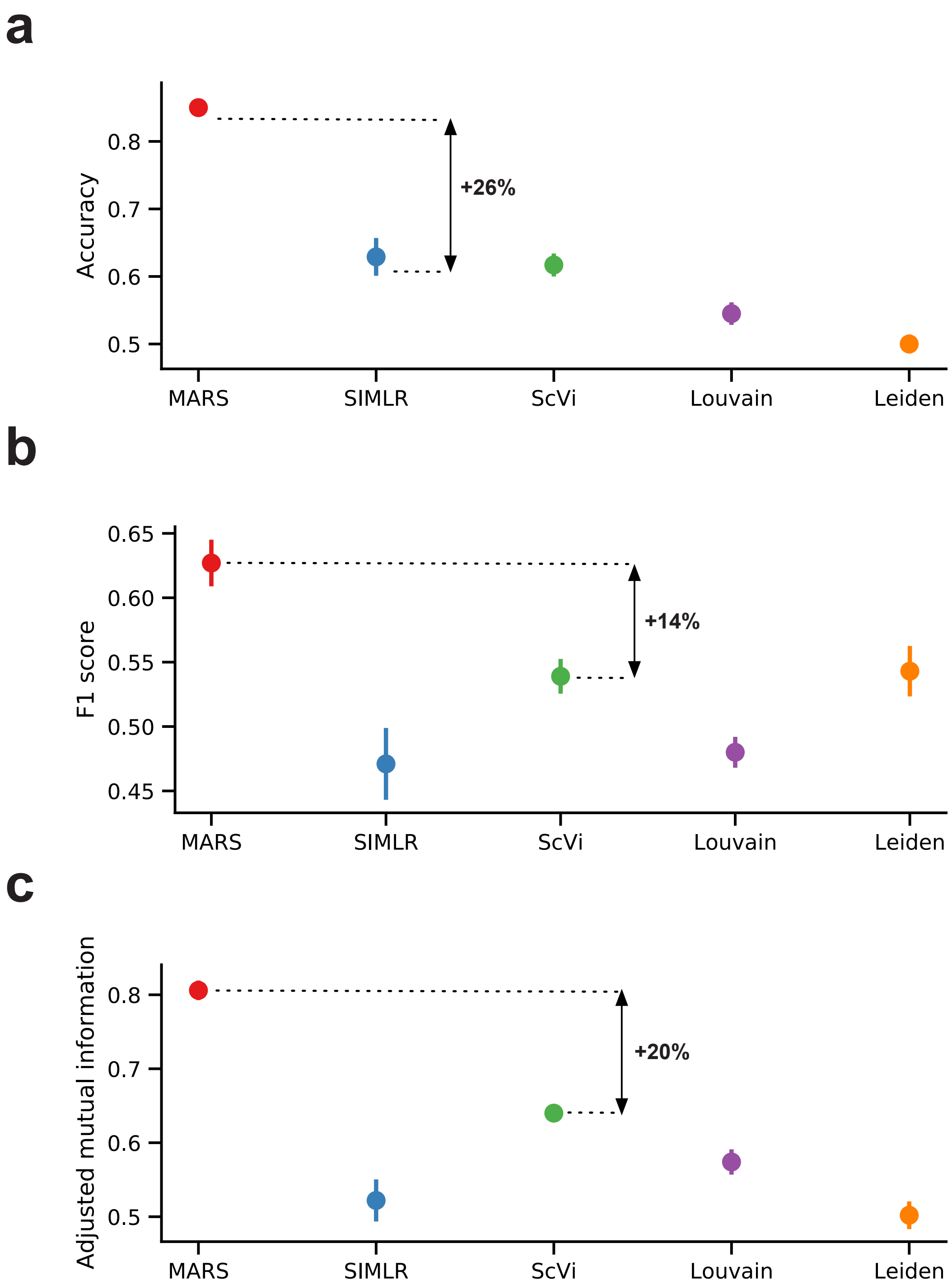


**Supplementary Figure 1:** Median performance of MARS and four baseline methods evaluated using **(a)** accuracy, **(b)** F1 score, and **(c)** adjusted mutual information. Baseline methods include SIMLR [6], ScVi [7], Louvain [4], and Leiden [5]. Median is calculated across 21 different tissues. For all measures, higher value indicates better performance. Error bars are standard errors estimated by bootstrapping cells within tissue with n=20 iterations. MARS is trained in leave-one-tissue-out manner, and the held out tissue was completely unannotated. MARS improves score of the second best performing method by 26%, 14%, and 20%, in terms of accuracy, F1 score, and adjusted mutual information, respectively.

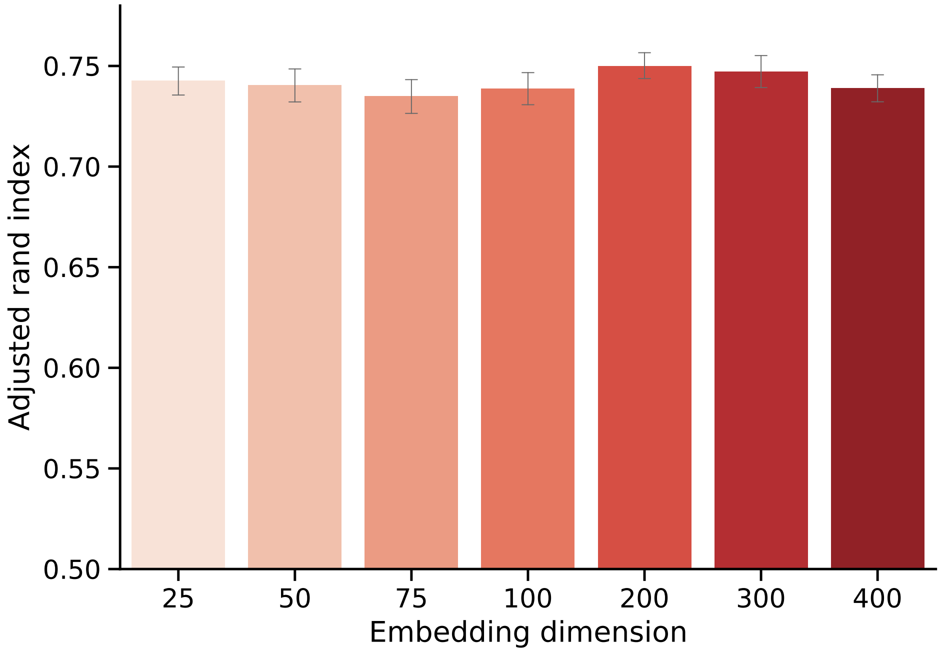


**Supplementary Figure 2:** Performance of MARS when varying number of neurons in the last layer of the neural network. Number of neurons in the last layer corresponds to the dimension of learned low-dimensional cell representation. Bars indicate average adjusted Rand index across $20$ runs of the method, where the value in each run corresponds to average score across $21$ tissues. Confidence intervals $(95\%)$ are determined using bootstrapping with $n = 1,000$ iterations. For each embedding dimension, we train MARS with all other parameters fixed.

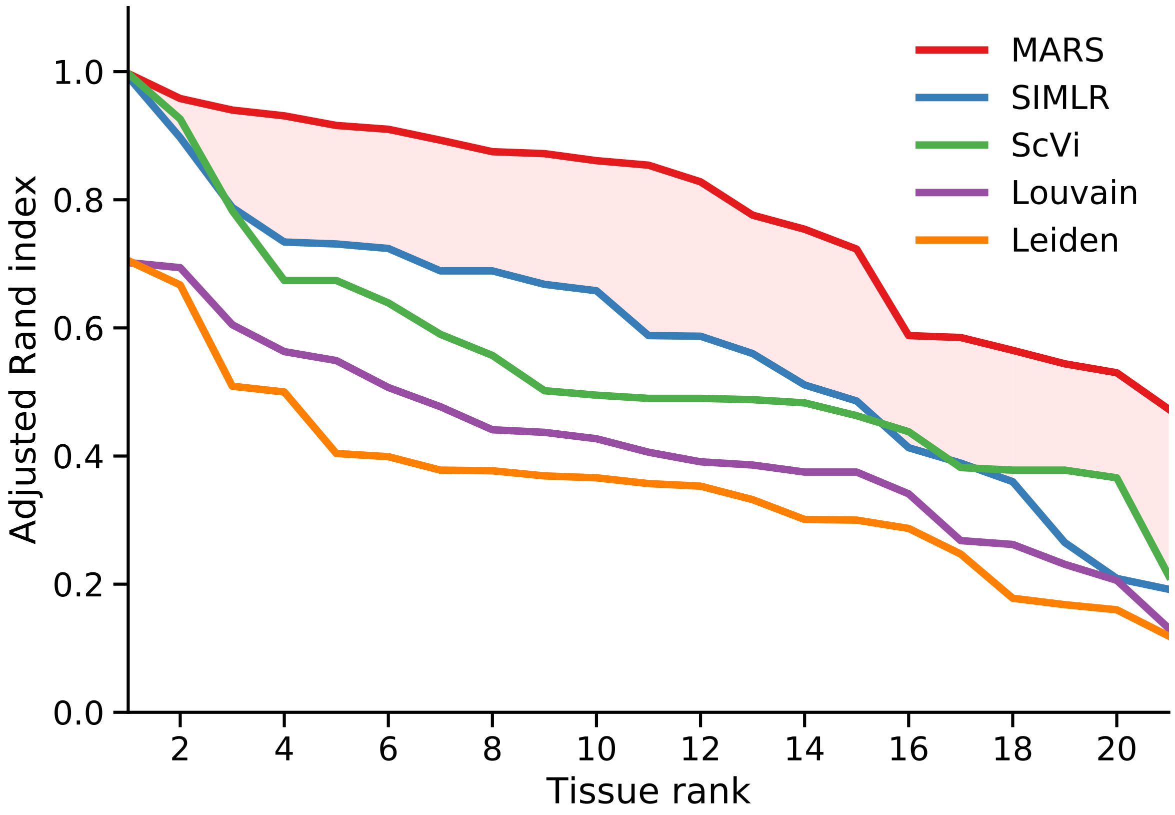


**Supplementary Figure 3:** Comparison of the MARS's clustering performance on individual tissues to baseline methods. Performance is measured as adjusted Rand index score. For each method, tissues are ranked in the decreasing order of the achieved score. Across all tissues, MARS achieves $26.1\%$ higher area under the curve compared to the SIMLR [6], and $30.1\%$ higher compared to the ScVi [7] baseline.

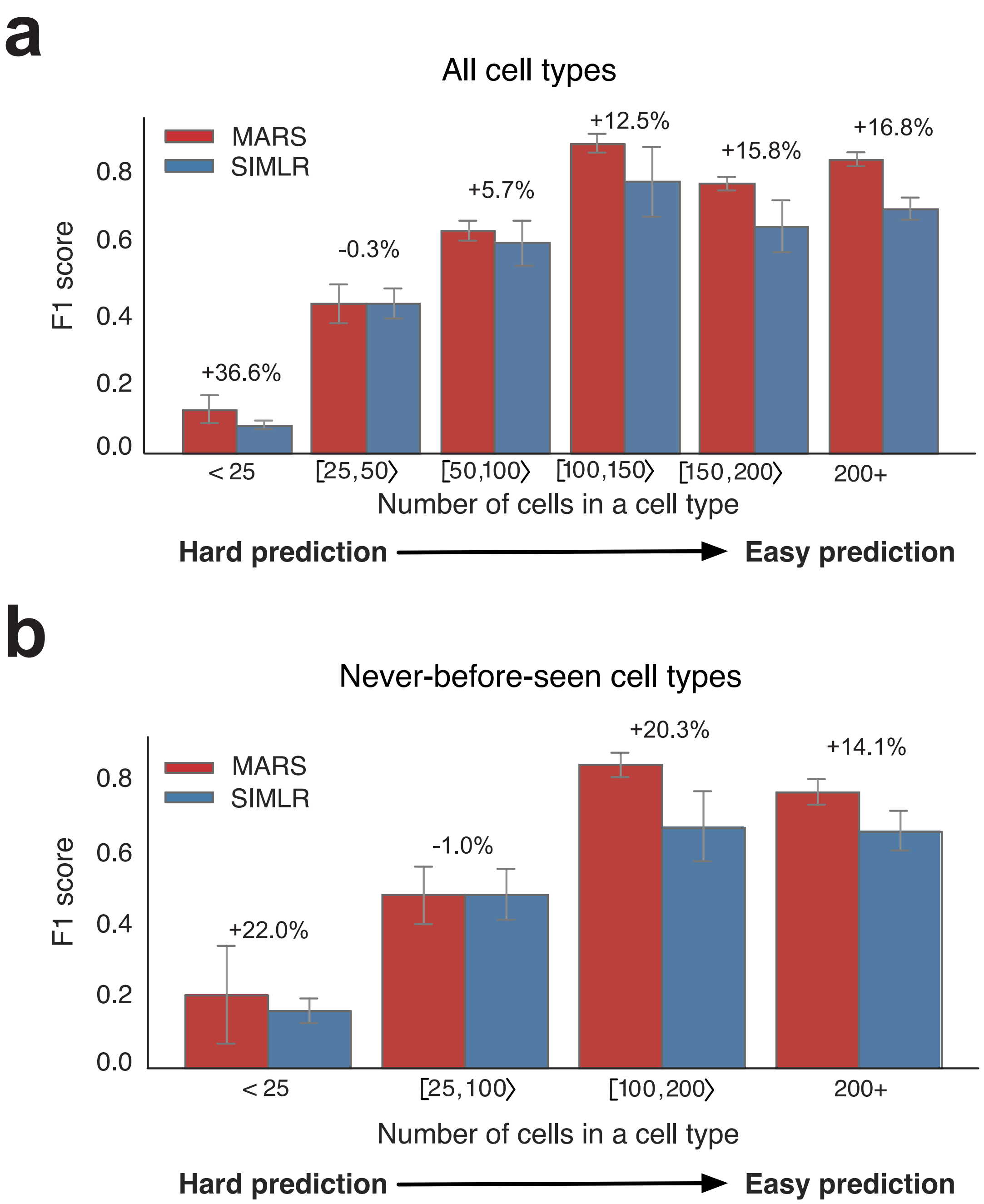


**Supplementary Figure 4:** **(a, b)** Cell-type level comparison of MARS's precision with the SIMLR on **(a)** all cell types, and **(b)** cell types that appear in only one tissue and have never been seen in the annotated experiment. Standard errors are estimated by bootstrapping cells within each tissue with n=20 iterations. Cell types are grouped based on the number of cells in the Tabula Muris annotations, where smaller number of cells means that it is harder to recognize cell type as a separate cluster.

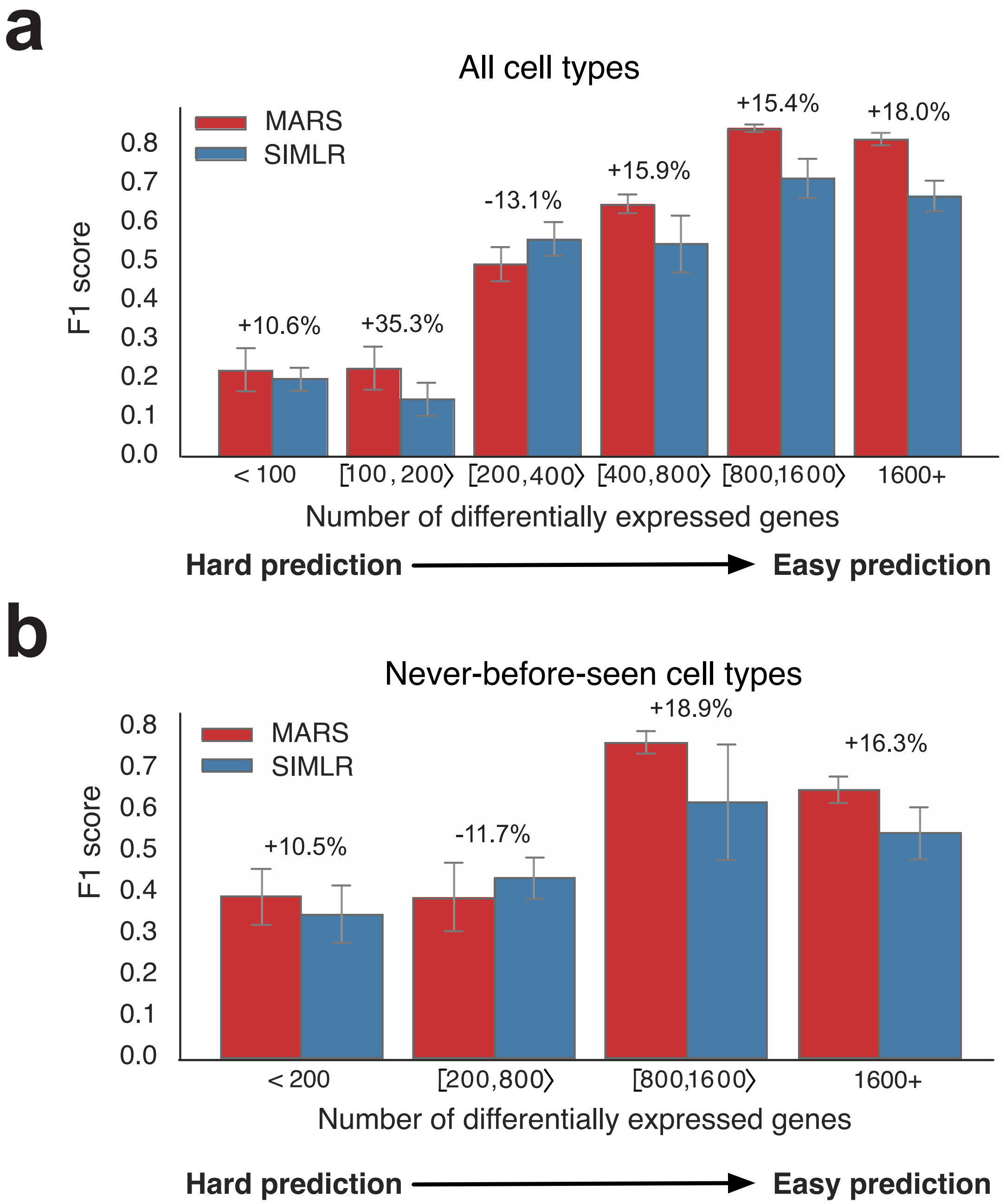


**Supplementary Figure 5: (a, b)** Cell-type level comparison of MARS's precision with the SIMLR on **(a)** all cell types, and **(b)** cell types that appear in only one tissue. Standard errors are estimated by bootstrapping cells within each tissue with n=20 iterations. Cell types are grouped based on the number of the differentially expressed genes calculated using the Tabula Muris annotations (Benjamini-Hochberg FDR adjusted p-value $< 0.01$; t-test).

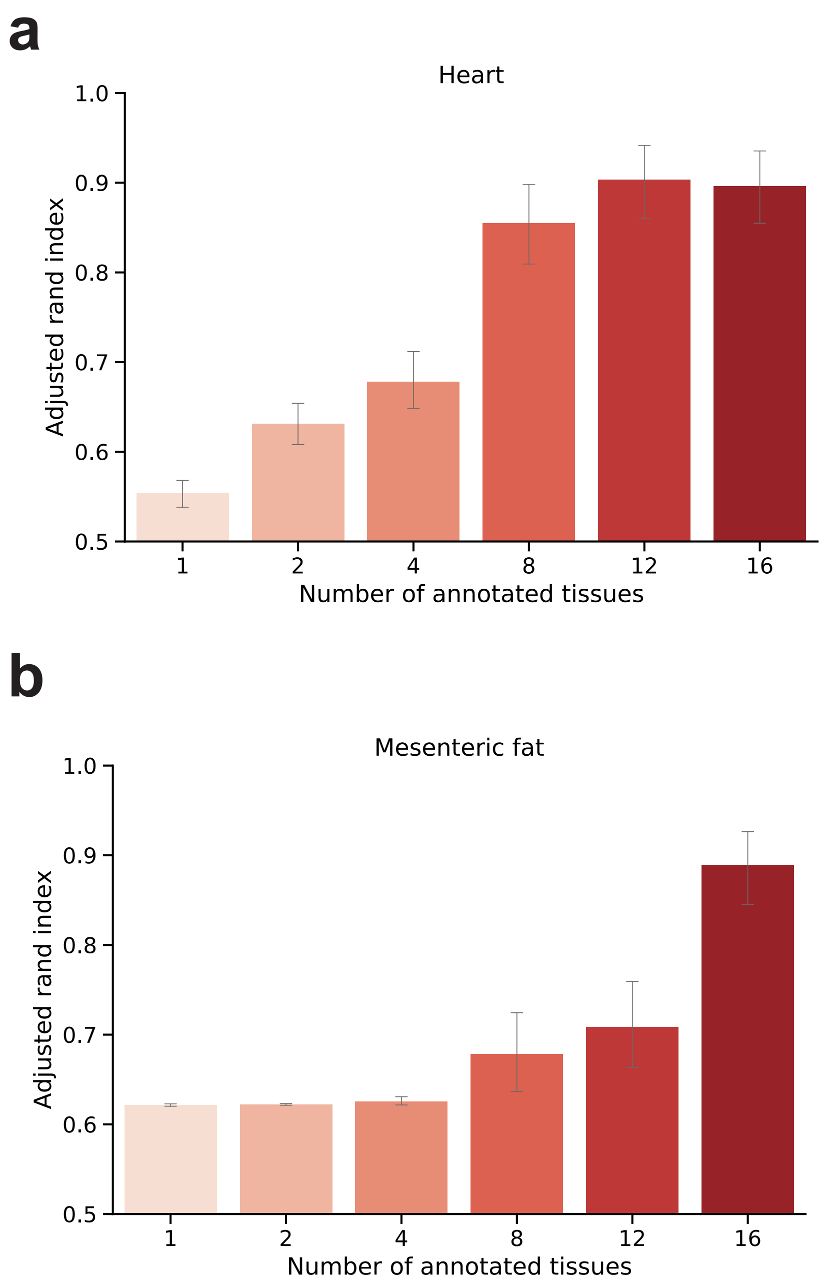


**Supplementary Figure 6:** **(a, b)** Effect of the number of annotated tissues in the meta-dataset on MARS's performance when using **(a)** heart tissue as unannotated experiment, and **(b)** mesenteric fat as unannotated experiment. Bars indicate average adjusted Rand index across $20$ runs of the method. Confidence intervals $(95\%)$ are determined using bootstrapping with $n = 1,000$ iterations. Annotated tissues are selected based on their similarity to an unannotated tissue, where similarity is computed as the euclidean distance of the mean gene expressions profiles between tissues. MARS improves performance when more tissues are included in the meta-dataset, implying that cross-tissue positive knowledge transfer is achieved even when tissues do not have similar gene expressions.

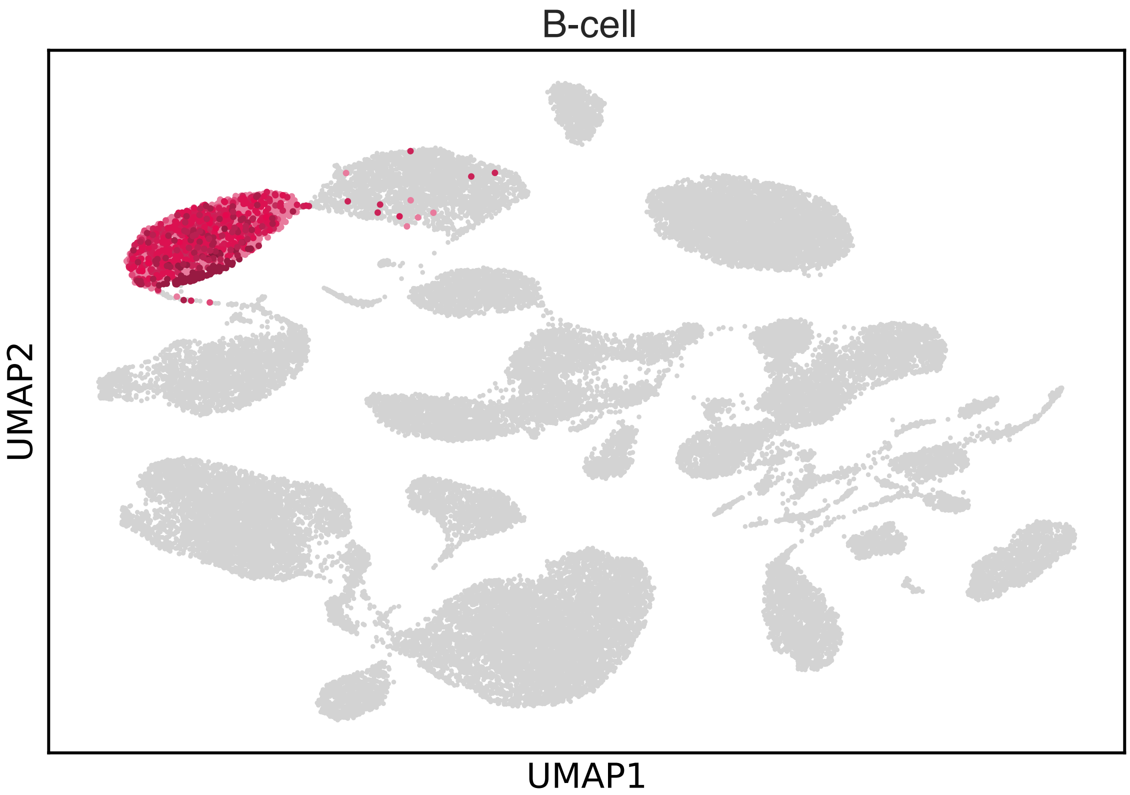


**Supplementary Figure 7:** Using MARS, B-cells in Tabula Muris data are extremely well aligned across 11 different tissues, including brown adipose tissue, diaphragm, gonodal fat, heart, kidney, limb muscle, lung, liver, mesenteric fat, subcutaneous fat, and spleen. Limb muscle is used as unnannotated tissue.
